# Suppression of transcytosis regulates zebrafish blood-brain barrier development

**DOI:** 10.1101/596221

**Authors:** Natasha M. O’Brown, Sean G. Megason, Chenghua Gu

## Abstract

As an optically transparent model organism with an endothelial blood-brain barrier (BBB), zebrafish offer a powerful tool to study the vertebrate BBB. However, the precise developmental profile of functional zebrafish BBB acquisition and the subcellular and molecular mechanisms governing the zebrafish BBB remain poorly characterized. Here we find a spatiotemporal gradient of barrier acquisition. Moreover, we capture the dynamics of developmental BBB leakage using live imaging, revealing a combination of steady accumulation in the parenchyma and sporadic bursts of tracer leakage. Electron microscopy studies further reveal that this steady accumulation results from high levels of transcytosis that are eventually suppressed, sealing the BBB. Finally, we demonstrate a key mammalian BBB regulator Mfsd2a, which inhibits transcytosis, plays a conserved role in zebrafish. *Mfsd2aa* mutants display increased larval and adult BBB permeability due to increased transcytosis. Our findings indicate a conserved developmental program of barrier acquisition between zebrafish and mice.

## Introduction

Blood vessels in the vertebrate brain are composed of a single layer of endothelial cells that possess distinct functional properties that allow the passage of necessary nutrients yet prevent unwanted entry of specific toxins and pathogens into the brain. This specialized endothelial layer forms the blood-brain barrier (BBB) and restricts the passage of substances between the blood and the brain parenchyma via two primary mechanisms: 1) specialized tight junction complexes between apposed endothelial cells to prevent intercellular transit (Reese and Karnovsky, 1967; Brightman and Reese, 1969) and 2) suppressing vesicular trafficking or transcytosis to prevent transcellular transit (Ben-Zvi et al., 2014; Andreone et al., 2015; Andreone et al., 2017). BBB selectivity is further refined with expression of substrate-specific transporters that dynamically regulate the influx of necessary nutrients and efflux of metabolic waste products (Sanchez-Covarrubias et al., 2014; Umans et al., 2017). While the BBB is comprised of endothelial cells, the surrounding perivascular cells including pericytes and astroglial cells, play a critical role in forming and maintaining barrier properties (Janzer and Raff, 1987; Armulik et al., 2010; Bell et al., 2010; Daneman et al., 2010; Wang et al., 2014). Collectively, endothelial cells and the surrounding perivascular cells form the neurovascular unit.

As the simplest genetic model organism with an endothelial BBB (Jeong et al., 2008), zebrafish offer a powerful tool to study the cellular and molecular properties of the vertebrate BBB (Xie et al., 2010; Vanhollebeke et al., 2015; Umans et al., 2017; O’Brown et al., 2018; Quiñonez-Silvero et al., 2019). Zebrafish have served as a great model system to study vascular biology due to their large clutch size, rapid and external development, and transparency for *in vivo* whole organism live-imaging (Lawson and Weinstein, 2002; Jin et al., 2005; Santoro et al., 2007; Armer et al., 2009; Herbert et al., 2009; Phng et al., 2009; Geudens et al., 2010; Herbert et al., 2012; Wilkinson and van Eeden, 2014; Franco et al., 2015; Vanhollebeke et al., 2015; Matsuoka et al., 2016; Ulrich et al., 2016; Galanternik et al., 2017; Stratman et al., 2017; Geudens et al., 2018). Additionally, with the advent of CRISPR-Cas9 technology, zebrafish provide an efficient genetic toolkit for targeted mutagenesis (Hwang et al., 2013; Gagnon et al., 2014; Ablain et al., 2015; Varshney et al., 2015; Albadri et al., 2017; Hogan and Schulte-Merker, 2017). However, the subcellular and molecular mechanisms governing the formation and maintenance of the zebrafish BBB remain poorly characterized. Expanding our understanding of the zebrafish BBB can thus reveal the mechanistic similarities between the zebrafish and mammalian BBB to further elevate the position of zebrafish as a model organism for studying the BBB.

Barrier properties of brain endothelial cells are induced by extrinsic signals from other cells in the surrounding microenvironment during development (Stewart and Wiley, 1981). In rodents, the BBB becomes functionally sealed in a spatiotemporal gradient, with the hindbrain and midbrain barriers becoming functional before the cortical barrier (Daneman et al., 2010; Ben-Zvi et al., 2014; Sohet et al., 2015). Within the cortex, barrier function is acquired along a ventral-lateral to dorsal-medial gradient (Ben-Zvi et al., 2014). In the zebrafish, existing studies have disagreed over the timing of zebrafish barrier formation, with some suggesting that BBB maturation occurs at 3 days post fertilization (dpf) (Jeong et al., 2008; Umans et al., 2017) and others providing a wide range beginning at 3 dpf and extending to 10 dpf (Fleming et al., 2013). These conflicting reports may be due to regional developmental gradients of barrier acquisition or differences in the experimental approaches used to assess BBB permeability such as the molecular weight of tracers and circulation time. To date, a thorough regional characterization of functional barrier acquisition has been lacking in zebrafish.

Recent work in the mammalian blood-retinal barrier has indicated that suppression of transcytosis governs functional barrier development (Chow and Gu, 2017). Interestingly endothelial cells at the leaky neonatal angiogenic front possess functional tight junction complexes halting the intercellular passage of the tracer protein Horseradish Peroxidase (HRP) at the so-called “kissing points”. In contrast, these endothelial cells exhibit high levels of HRP-filled vesicles compared to functionally sealed proximal vessels. Moreover, these areas of elevated vesicular trafficking continue to correspond with barrier permeability at the angiogenic front until the barrier seals (Chow and Gu, 2017). Work in the mouse BBB has also demonstrated the importance of suppressing transcytosis in determining barrier permeability. Mice lacking the major facilitator super family domain containing 2a (Mfsd2a) lipid transporter exhibit increased levels of caveolae vesicles in CNS endothelial cells, resulting in increased barrier permeability (Ben-Zvi et al., 2014; Andreone et al., 2017). The subcellular and molecular mechanisms of zebrafish BBB acquisition have yet to be elucidated.

Here in zebrafish, we find a spatiotemporal gradient of barrier acquisition, and capture the dynamics of developmental BBB leakage using time lapse live imaging. We further find a conserved role for transcytosis suppression in determining barrier properties, both during normal development and in *mfsd2aa* mutants.

## Results

### Posterior-Anterior Gradient of Zebrafish BBB Development

To determine when and how the zebrafish BBB becomes functional in different brain regions, we performed intracardiac injections of fluorescently-conjugated tracers (1 kDa NHS and 10 kDa Dextran) simultaneously at different developmental stages and imaged live fish after one hour of tracer circulation (Figure 1A and 1B). We used a combination of different molecular weight tracers to tease apart potential avenues of leakage, as tight junctional defects result specifically in the leakage of low molecular weight tracers 1 kDa and below into the brain parenchyma (Nitta et al., 2003; Campbell et al. 2008; Sohet et al., 2015; Yanagida et al. 2017). At 3 dpf, we observed a sealed barrier in the hindbrain as previously described (Jeong et al., 2008), with only a few of the parenchymal cells taking up the injected tracer (average of 2 ± 0.3 cells/embryo with NHS and 2 ± 0.4 cells/embryo with Dextran; Figure 1 – Supplement 1), which we quantify as a proxy of tracer leakage into the brain. However, in the midbrain we observed an increased number of parenchymal cells that accumulated the circulating tracers (average of 24 ± 1 cells/embryo with NHS and 24 ± 1 cells/embryo with Dextran; Figure 1C and 1D), indicating that the tracers leaked out of the blood vessels into the brain and that the midbrain barrier is not yet functional. In addition to the use of exogenous injected fluorescent tracers, we also assayed BBB permeability with an endogenous transgenic serum DBP-EGFP fusion protein (Tg(*l-fabp:DBP-EGFP*)) to account for injection artifacts (Xie et al., 2010). At 3 dpf, we observed similar leakage patterns with the transgenic serum protein as we did with the injected tracers (average of 24 ± 1 cells/embryo in the midbrain and average of 2 ± 0.4 cells/embryo in the hindbrain; Figure 1C and D; Figure 1 – Supplement 1). At 4 dpf, the BBB in the hindbrain is completely functional with few tracer-filled parenchymal cells (average of 3 ± 0.4 cells/embryo with NHS, 2 ± 0.4 cells/embryo with Dextran and DBP-EGFP; Figure 1 – Supplement 1). However, the midbrain BBB remains leaky (average of 23 ± 1 cells/embryo with NHS and DBP-EGFP and 24 ± 1 cells/embryo with Dextran; p=0.68 compared to 3 dpf, one-way ANOVA; Figure 1C and 1D). However, at 5 dpf the number of midbrain parenchymal cells that uptake the tracers is dramatically reduced (average of 9 ± 1 cells/embryo with NHS, Dextran, and DBP-EGFP; p<0.0001, one-way ANOVA; Figure 1B and 1C) and no change was observed in the hindbrain (Figure 1 – Supplement 1). No significant change was observed in midbrain permeability from 5 to 6 dpf (average of 8 ± 1 cells/embryo with NHS, Dextran, and DBP-EGFP; p=0.904, one-way ANOVA; Figure 1C and 1D), indicating that the midbrain barrier becomes sealed at 5 dpf. We did not assay the forebrain at any of these stages due to the fact that it remains avascularized until 5 dpf. Of note, all three tracers showed nearly indistinguishable patterns of uptake, with similar numbers of cells and many of the same cells simultaneously taking up both injected tracers (Figure 1; Figure 1 – Supplement), suggesting that the leakage is not due to tight junctional defects but may rather be due to an increase in vesicular trafficking.

**Figure 1.**
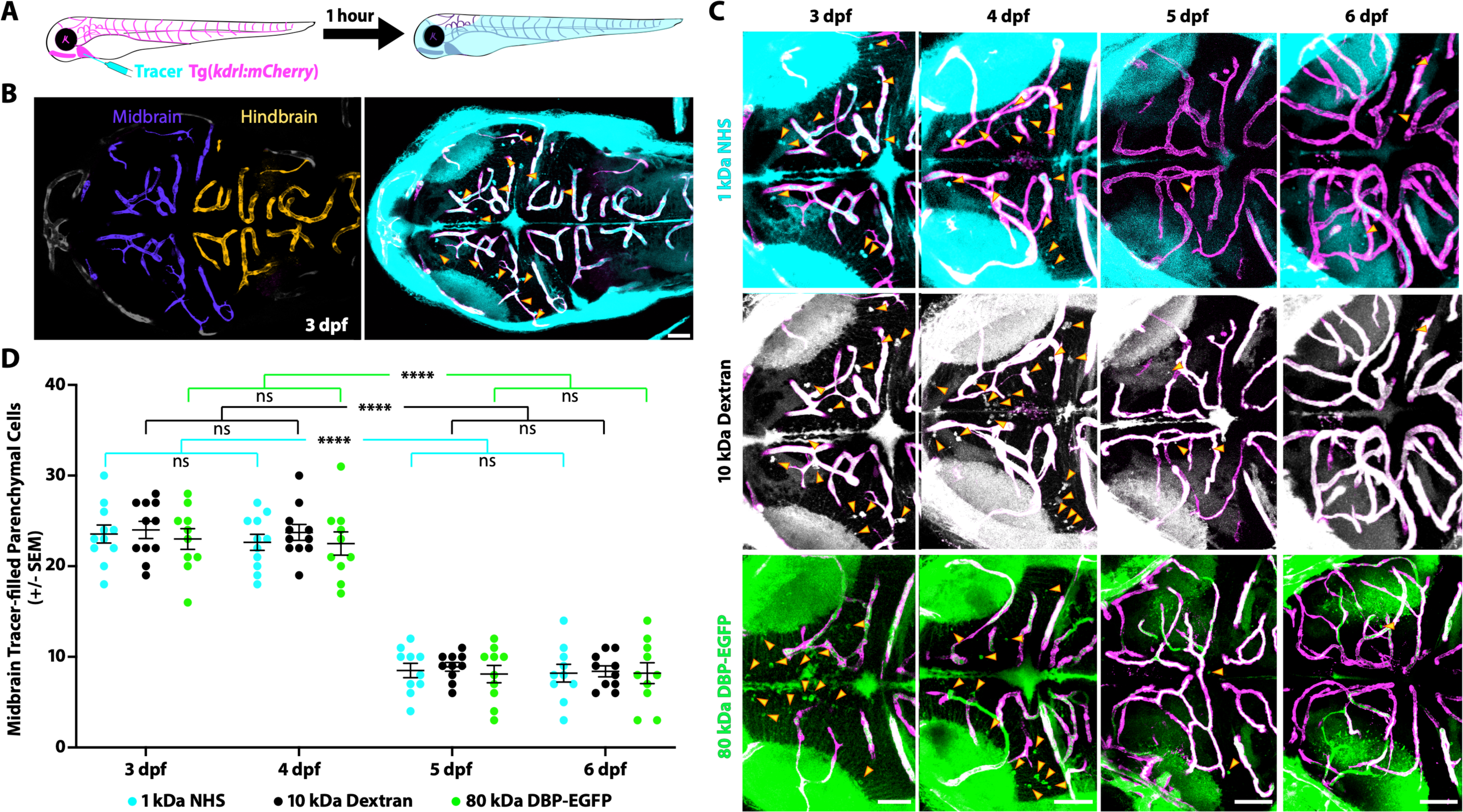
The midbrain BBB becomes functional at 5 dpf. (A) Diagram of the tracer leakage assay. Fluorescently conjugated tracers (turquoise) were injected intracardially into transgenic fish that express mCherry in the vasculature (magenta; Tg(*kdrl:mCherry*)) and allowed to circulate for one hour before imaging. (B) Dorsal view maximum intensity projection of the larval brain vasculature at 3 dpf. Left image is pseudo-colored to demarcate the midbrain (violet) and the hindbrain (gold) vasculature. Right image shows the NHS tracer (turquoise) in the entire larval brain, with a large number of tracer-filled parenchymal cells marked by yellow arrowheads in the midbrain. (C) Representative dorsal view maximum intensity projections of larval zebrafish midbrains at different developmental stages reveal increased permeability at 3 and 4 dpf compared to 5 and 6 dpf. The increased early permeability was observed with two injected tracers of different sizes, a 1 kDa NHS (turquoise) and a 10 kDa Dextran (white), as well with an 80 kDa transgenic serum protein DBP-EGFP (green). Yellow arrowheads demarcate the tracer-filled parenchymal cells outside of the vasculature (magenta). Scale bars represent 50 µm. (D) Quantification of tracer-filled parenchymal cells in the midbrain between 3 and 6 dpf reveals a significant decrease in tracer uptake at 5 dpf. There was no difference in the number of parenchymal cells that picked up the different tracers at any time point. There was no significant change from 3 to 4 dpf or from 5 to 6 dpf, suggesting that the barrier seals around 5 dpf. N = 10-11 fish, each represented as a single dot on the plot. The mean and the standard error are drawn in black for each tracer and stage. **** p<0.0001, ns is not significant.

### Time lapse Imaging Reveals Two Modes of Developmental Leakage

The vast majority of studies interrogating BBB permeability have relied on observing leakage in fixed static images from mutant mice, which does not reveal the dynamic nature of the BBB. A huge advantage of zebrafish is the ability to examine biological processes *in vivo* in real-time, providing us with the unique opportunity to observe the process of BBB maturation (Figure 1). To examine the developmental dynamics of barrier leakage, we injected embryos at 3 dpf with fluorescently-labeled 10 kDa Dextran and performed time lapse live imaging for an hour following injection, with time 0 being an average of 8 minutes post-injection. We observed a gradual and diffuse increase in extravascular Dextran intensity over time within the brain parenchyma (slope of 1.301e-4 intensity/sec; Videos 1 and 2; Figure 2A, 2C and 2E). In addition to the time-dependent increase in overall Dextran intensity in the brain parenchyma, we also observed parenchymal cells taking up tracer, both directly adjacent to blood vessels and at a small distance away (Figure 2A and 2C), as in the static images of tracer leakage during development (Figure 1). Interestingly, at 3 dpf we observed an actively sprouting cell that extended away from the established vessel in a sporadic fashion (Video 2; Figure 2A). After twenty minutes of rambling migration, this sprout suddenly released a large bolus of Dextran that appeared to be taken up by a single parenchymal cell. Two minutes later, this same sprout released a second bolus of Dextran on the opposite side (Video 2). These rare large bursts of leakage were not unique to this sprout; they were also sporadically observed from established blood vessels (Video 1; Figure 2A). This data revealed two types of leakage occurring during this early developmental stage: steady and diffuse Dextran leakage into the parenchyma that makes up the vast majority of observed leakage and rare large bursts of leakage. When we performed these same time lapse experiments at 5 dpf, we observed significantly less overall tracer accumulation in the brain parenchyma over the course of the hour in addition to reduced rates of tracer accumulation (slope of 4.641e-5 intensity/sec; Video 3; p<0.001, Mann-Whitney test; Figure 2B, 2D and 2E). Furthermore, 5 dpf fish never exhibited bursts of tracer leakage as observed at 3 dpf. Overall, 5 dpf fish had far fewer parenchymal cells filling with Dextran than at 3 dpf, as observed in the static tracer leakage assays (Figure 1). We interpret these data to suggest that at early stages, most tracer leakage occurs broadly through vessel walls potentially via unrestricted transcytosis with an occasional transient rupture to endothelial integrity that lead to these large bursts of leakage. Once tracer is leaked into the intercellular space of the parenchyma it is taken up by select parenchymal cells. As the BBB matures both sources of leakage sharply decrease.

**Figure 2.**
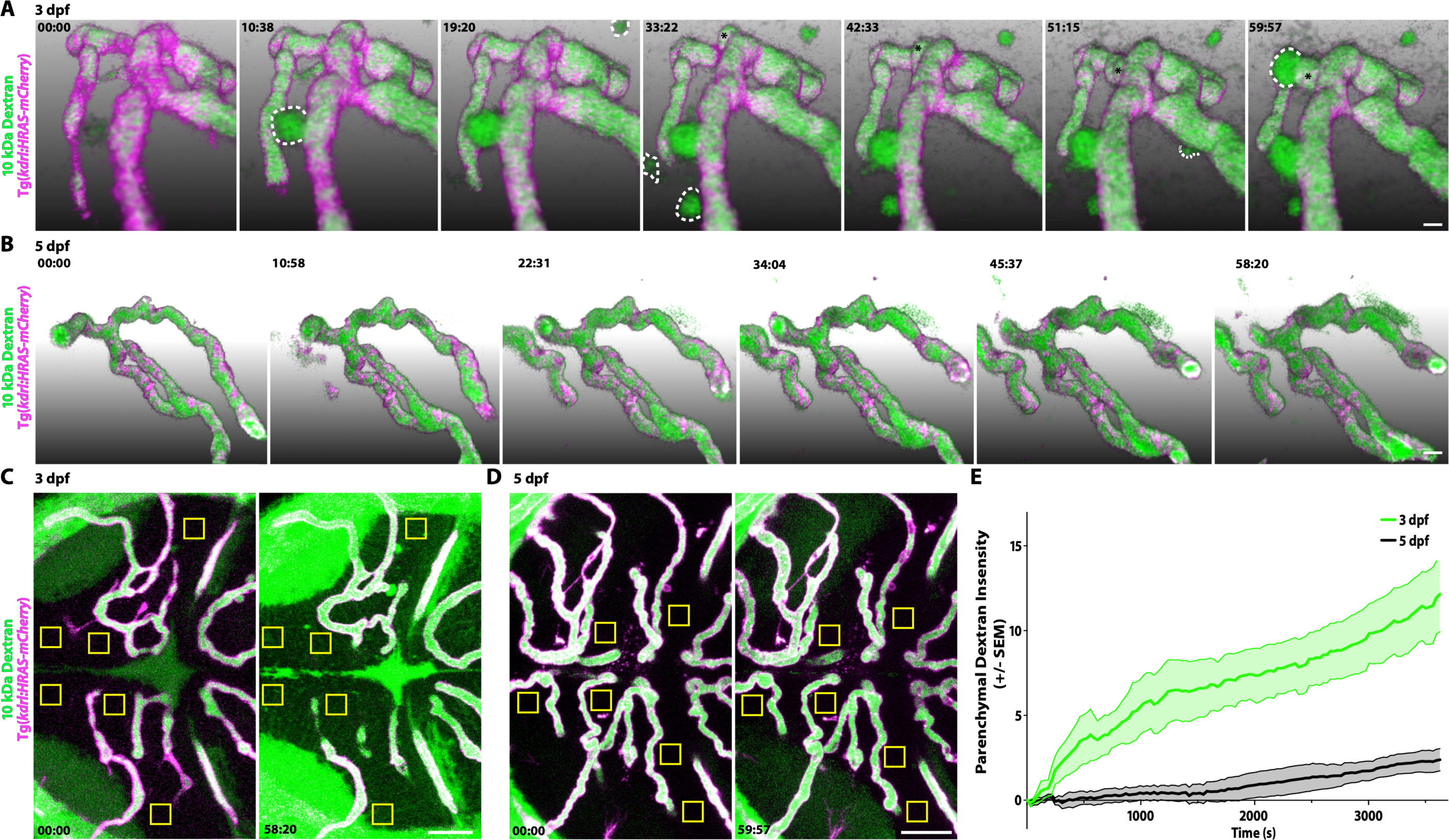
Dynamic tracer leakage in the developing BBB via live imaging. (A) Time course stills from Video 1 of tracer leakage at 3 dpf reveal an increase in parenchymal cells absorbing the Dextran tracer (outlined by dashed white lines) as well as a general increase in overall Dextran (green) intensity outside of the vasculature (magenta). An angiogenic tip cell becomes apparent at 33:23 and is demarcated by an asterisk (*). This tip cell produces two separate bursts of leakage observed in Video 2. (B) Time course stills of Dextran tracer dynamics at 5 dpf reveals a mature BBB, with reduced overall Dextran extravasation into the brain parenchyma. The scale bars represent 10 µm. (C and D) Dorsal maximum intensity projection of the midbrain at 3 dpf (C) and 5 dpf (D) at the first and last time point examined. While there is a large increase in overall parenchymal Dextran intensity over time at 3 dpf, the 5 dpf midbrain parenchyma appears relatively unaltered after an hour of Dextran circulation. Boxed regions are representative of the six areas per fish used for analysis in E. The scale bars represent 50 µm. (E) Quantification of Dextran intensity in the brain parenchyma over time at 3 dpf (green) and 5 dpf (black) shows a significant difference in tracer leakage dynamics (p<0.0001, Mann Whitney U test), with both more total Dextran accumulation and a faster rate of Dextran accumulation in the brain parenchyma at 3 dpf. N=6 fish with 6 regions analyzed per fish.

### Suppression of Transcytosis Governs BBB Development

Given the observed differences in permeability during larval development, we next sought to determine the subcellular mechanisms underlying the development of a functional BBB. Therefore, we assessed BBB properties by performing intracardiac injections of electron-dense NHS-gold nanoparticles (5 nm) followed by transmission electron microscopy (TEM) at different developmental stages when the midbrain barrier is leaky (3 dpf) and when it is functionally sealed to fluorescent tracers (5 and 7 dpf; Figure 3). All of the blood vessels analyzed had a maximal diameter of 5 µm to enrich for capillaries and small veins. At 3 dpf, we found blood vessels in direct contact with neurons and pericytes, with the pericytes sharing the endothelial basement membrane (Figure 3A), as in the mammalian neurovascular unit. During this leaky developmental stage, the endothelial basement membrane was filled with the electron-dense gold nanoparticles (Figure 3A and 3B; turquoise arrowheads) with an average gold intensity of 1.29 ± 0.07 (luminal gold intensity normalized to 1.0; Figure 3G), further demonstrating that the BBB is immature at 3 dpf. To decipher how the gold nanoparticles traverse from the lumen to the basement membrane, we first looked at the tight junctions to see if the nanoparticles passed between apposed endothelial cells. After careful examination, we observed that the nanoparticles were halted at the “kissing points” between endothelial cells, indicating that tight junctions were functional prior to formation of a functional BBB (49/49 functional tight junctions; Figure 3B; green arrowhead). As transcytosis has been implicated in the maturation of the blood-retinal barrier (Chow and Gu, 2017), we next examined the levels of luminal and abluminal flask-shaped vesicles filled with gold nanoparticles as a means of assessing transcytosis. Quantification of gold-filled luminal and abluminal flask-shaped vesicles revealed an average of 0.21 ± 0.03 and 0.17 ± 0.02 vesicles/µm, respectively (Figure 3H and 3I). These data reveal that the basement membrane becomes filled with gold nanoparticles via vesicular transport rather than intercellular passage between immature tight junctions.

**Figure 3.**
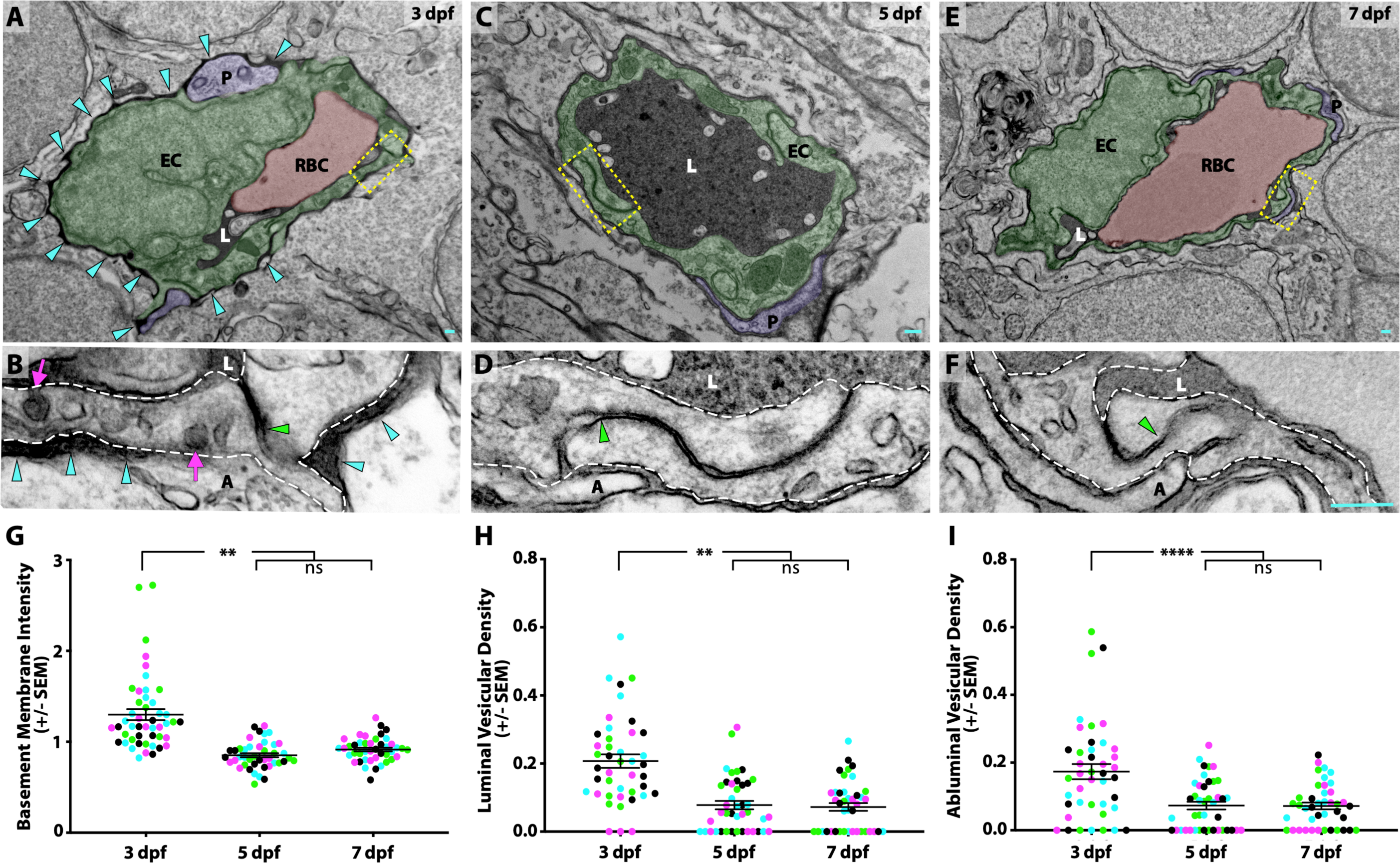
Suppression of transcytosis determines the timing of functional BBB formation. (A, C, E) TEM images of individual blood vessel cross-sections after injection of electron-dense gold nanoparticles at 3 dpf (A), 5 dpf (C), and 7 dpf (E). Endothelial cells (EC) are pseudo-colored green, pericytes (P) are pseudo-colored purple and red blood cells (RBC) are pseudo-colored red when present in the lumen (L). Turquoise arrowheads highlight the gold-filled basement membrane at 3 dpf (A). (B, D, F) High magnification images (25000x) of the areas boxed in A, C, and E, respectively, with the endothelial cells outlined with white dashed lines. The images are oriented with the lumen (L) on top and the ablumen (A) on the bottom. Tight junctions are functional as early as 3 dpf (B), as seen by their ability to halt the gold nanoparticles at the so-called “kissing point” (green arrowhead), and remain functional throughout development (D and F). Even though the tight junctions are functional at 3 dpf, the endothelial basement membrane is filled with electron-dense gold nanoparticles (B, turquoise arrowheads). This appears to be due to an elevated level of luminal and abluminal gold-filled vesicles (magenta arrows). The scale bars represent 200 nm. (G) Quantification of the endothelial basement membrane gold intensity normalized to luminal gold intensity. (H and I) Quantification of the vesicular densities both on the luminal (H) and abluminal (I) membrane of endothelial cells reveals a suppression of vesicular densities beginning at 5 dpf that remains constant at 7 dpf. N=4 fish, each marked with a different color, with at least 10 blood vessels quantified for each fish. **** p<0.0001, ** p<0.01, ns is not significant.

When we repeated this assay at 5 dpf, when the barrier becomes less permeable to fluorescently conjugated tracers (Figures 1 and 2), we still observe close contacts between endothelial cells and pericytes (Figure 3C). At 5 dpf, the basement membrane was noticeably lacking gold particles as compared to 3 dpf (Figure 3D), with a reduced average gold intensity of 0.85 ± 0.02 (Figure 3G). Like at 3 dpf, the tight junctions remained functional at 5 dpf, based on their capacity to halt gold nanoparticles at the kissing points between neighboring endothelial cells (76/76 functional tight junctions; Figure 3D). However, the levels of vesicles both luminally and abluminally were notably decreased to 0.08 ± 0.01 and 0.07 ± 0.01 vesicles/µm, respectively (p<0.001 (luminal) and p<0.0001 (abluminal), nested one-way ANOVA; Figure 3H and 3I). At 7 dpf the neurovascular cellular interactions remained constant with endothelial cells in close contact with pericytes and neurons, and with an unfilled basement membrane (average gold intensity of 0.91 ± 0.02; Figure 3E and 3G). Importantly, the tight junctions remained functional (51/51 functional tight junctions), and the low vesicular densities observed at 5 dpf remained comparably low at 7 dpf with 0.07 ± 0.01 vesicles/µm, both luminally and abluminally (Figure 3F, 3H and 3I). Taken together with the fluorescent tracer data, our data suggests that the zebrafish BBB becomes functional at 5 dpf via suppression of vesicular trafficking.

### Conserved Role of Mfsd2a in Determining BBB Function

Given the important role of suppressing transcytosis in determining the developmental maturation of the zebrafish BBB, we wondered whether the key mammalian barrier regulator Mfsd2a, which suppresses caveolae mediated transcytosis (Ben-Zvi et al., 2014; Andreone et al., 2017), plays a conserved role in zebrafish. Zebrafish contain two paralogues of *Mfsd2a*, *mfsd2aa* and *mfsd2ab*. *Mfsd2aa* is 61% identical to human *MFSD2A* and 62% identical to mouse *Mfsd2a* (Figure 4 – Supplement 1A). *Mfsd2ab*, on the other hand, is 64% identical to human *MFSD2A* and mouse *Mfsd2a* (Figure 4 – Supplement 1A). The two paralogues are only 68% identical to each other, but they both contain the lipid binding domain that is critical for governing barrier properties (Figure 4 – Supplement 1A; Andreone et al., 2017). Given the lack of a clear paralogue that most closely resembles *Mfsd2a*, we generated CRISPR mutants for both paralogues independently. *Mfsd2aa^hm37/hm37^*mutants have a 7 bp deletion in exon 2 (Figure 4 – Supplement 1B) that is predicted to lead to a premature stop codon at amino acid 82 (Figure 4 – Supplement 1A; black box). Homozygous *mfsd2aa* mutants are viable and fertile. *Mfsd2ab^hm38/hm38^* mutants have a 19 bp deletion in exon 5 (Figure 4 – Supplement 1B) that is predicted to lead to a premature stop codon at amino acid 175 (Figure 4 – Supplement 1A; black box). Homozygous *mfsd2ab* mutants are also viable and fertile. Neither mutant displayed obvious angiogenic defects, similar to the normal vasculature observed in mouse *Mfsd2a* mutants (Ben-Zvi et al., 2014).

**Figure 4.**
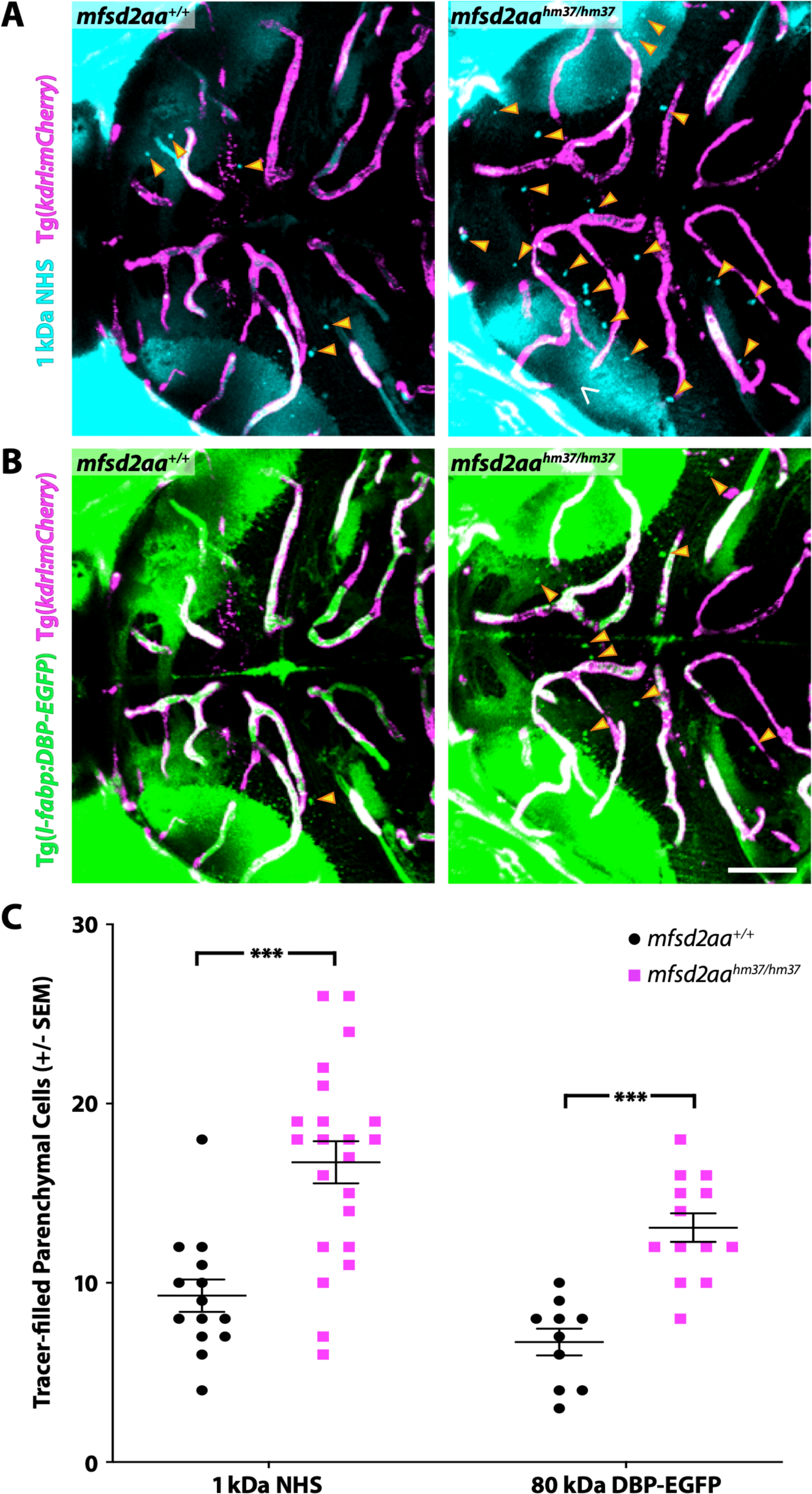
*Mfsd2aa* mutants exhibit increased BBB permeability. (A) Representative maximum intensity projection images of the midbrain of wildtype and *mfsd2aa* mutants injected with a fluorescent 1 kDa NHS tracer (turquoise) at 5 dpf. *Mfsd2aa* mutants have an increased number of NHS-filled parenchymal cells (yellow arrowheads) that lie outside of the vasculature (magenta; Tg(*kdrl:mCherry*)). (B) Representative maximum intensity projection images of the midbrain of wildtype and *mfsd2aa* mutants expressing the fluorescently labelled 80 kDa transgenic serum protein DBP-EGFP (green) at 5 dpf. *Mfsd2aa* mutants have an increased number of DBP-EGFP-filled parenchymal cells (yellow arrowheads) compared to wildtype siblings. The scale bar represents 50 µm. (C) Quantification of tracer-filled (NHS and DBP-EGFP) parenchymal cells in the midbrain of wildtype (black) and *mfsd2aa* mutants (magenta). *Mfsd2aa* mutants have a significantly increased number of tracer-filled parenchymal cells, both for the injected NHS (A) and the endogenous transgene DBP-EGFP (B). Each individual fish measured is displayed as a single point. The mean and the standard error are drawn as black lines. *** p < 0.001, ** p < 0.01.

To investigate whether either paralogue was necessary for barrier formation, we performed NHS tracer injection assays as we did to identify the developmental timeline of barrier formation and assayed for midbrain leakage at 5 dpf. In addition to the use of the exogenous injected fluorescent tracer (1 kDa NHS), we also assayed the leakage of the endogenous transgenic serum DBP-EGFP fusion protein (Tg(*l-fabp:DBP-EGFP*)). *Mfsd2aa* mutants displayed about a two-fold increase in midbrain tracer-containing parenchymal cells, both for the injected 1 kDa NHS and the endogenous serum transgene 80 kDa DBP-EGFP, at 5 dpf compared to wildtype sibling controls (Figure 4). These data suggest that mfsd2aa plays a similar role to mouse Mfsd2a in determining barrier properties. Conversely, *mfsd2ab* mutants displayed similarly low levels of tracer-containing parenchymal cells to wildtype controls at 5 dpf in the midbrain (Figure 4 – Supplement 2), indicating that mfsd2ab is dispensable for functional barrier formation. To see if the loss of both paralogues resulted in increased barrier permeability, we investigated *mfsd2aa^hm37/hm37^; mfsd2ab^hm38/hm38^*double mutants for BBB permeability using the endogenous serum DBP-EGFP fusion protein (Figure 4 – Supplement 3). While *mfsd2aa* single mutants displayed the previously observed increase in barrier permeability and *mfsd2ab* single mutants did not, *mfsd2aa^hm37/hm37^; mfsd2ab^hm38/hm38^* double mutants displayed similar levels of increased barrier permeability to the *mfsd2aa* single mutants (Figure 4 – Supplement 3). Taken together, these data suggest that the two paralogues, while structurally similar, play different roles, with zebrafish mfsd2aa and mammalian Mfsd2a sharing a conserved role in determining barrier permeability.

Given the leakage phenotype in *mfsd2aa* mutants at 5 dpf, we next wanted to examine whether the leakage phenotype persisted into adulthood. To address this, we performed retro-orbital injections of HRP, which has been shown to be confined within the adult zebrafish brain vasculature (Jeong et al., 2008), and allowed the HRP to circulate for 30 minutes. As expected, the wildtype siblings retained the HRP within their blood vessels (Figure 5A). However, *mfsd2aa* mutants exhibited HRP extravasation into the brain parenchyma (Figure 5B), suggesting that the leakage phenotype was not limited to larval fish. Finally, to determine whether this increased permeability was due to increased transcytosis as in *Mfsd2a* knockout mice, we measured vesicular density in capillaries with luminal diameters less than 5 µm from adult mutant and wildtype siblings using TEM. The *mfsd2aa* mutant blood vessels appeared morphologically normal by TEM, composed of a thin single layer of endothelial cells in close contact with pericytes, as observed in their wildtype siblings (Figure 5C and 5D). A closer examination of the endothelial cells revealed electron-dense tight junction complexes between all apposed endothelial cells in both wildtype and *mfsd2aa* mutant fish (Figure 5E and 5F). However, while wildtype fish display similarly low levels of luminal (0.1 vesicles/µm) and abluminal (0.11 vesicles/µm) vesicular densities to those observed at 7 dpf (Figure 5E-5H), *mfsd2aa* mutant fish display a significant increase in luminal and abluminal vesicular densities (0.29 and 0.27 vesicles/µm, respectively) compared to wildtype siblings (Figure 5E-5H). Interestingly this increase in vesicular abundance is even higher than that observed during early barrier development at 3 dpf (Figure 2). These data also suggest that the increased tracer leakage observed in *mfsd2aa* mutants results from an increase in vesicular trafficking across the BBB, further supporting a conserved role for mfsd2aa in determining barrier properties. In contrast to *mfsd2aa* mutants, *mfsd2ab* mutants displayed similar levels of abluminal and luminal vesicular pit density to wildtype siblings (Figure 5 – Supplement 1), further demonstrating that the paralog *mfsd2ab* does not play a conserved role in determining barrier properties in zebrafish.

**Figure 5.**
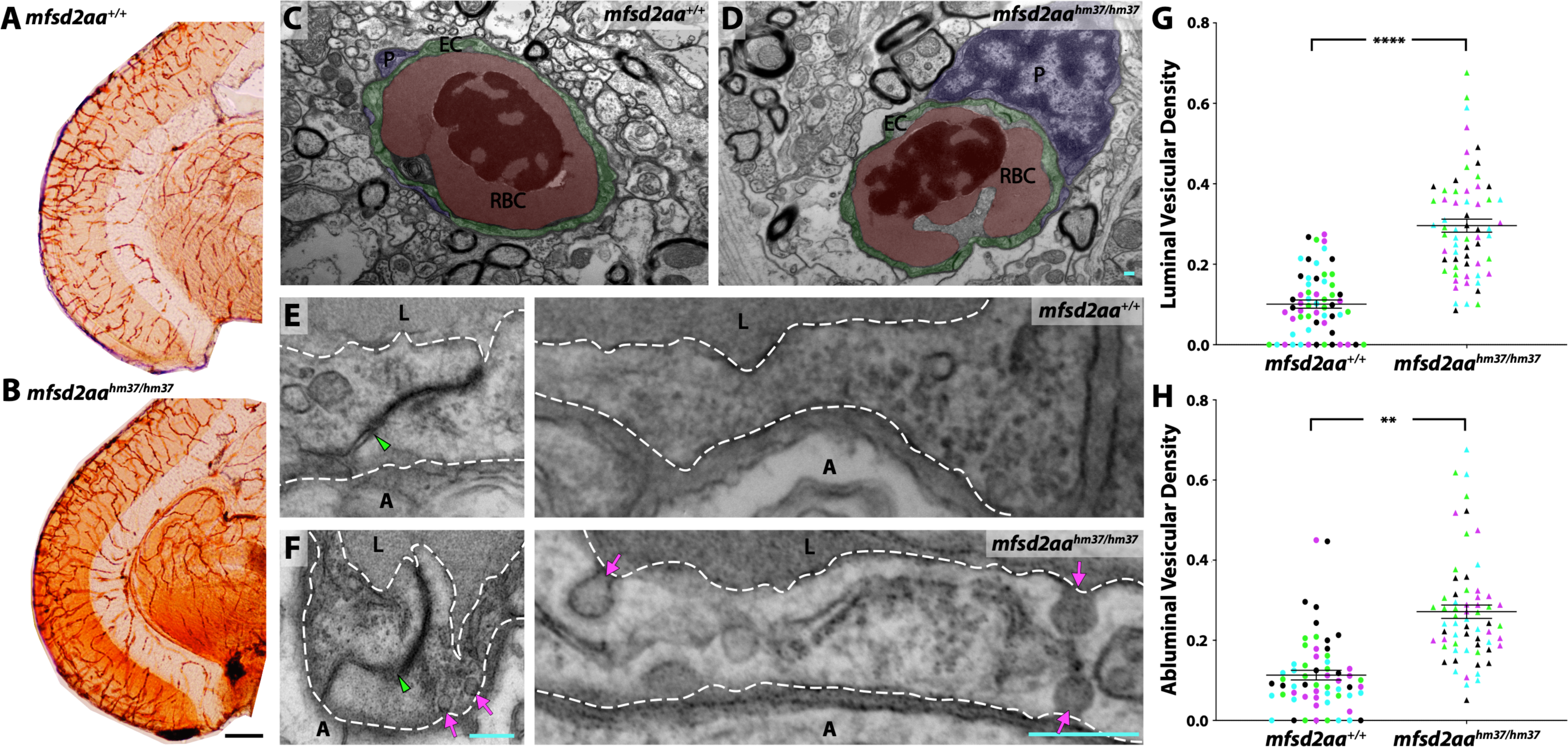
M*fsd2aa* mutants exhibit increased transcytosis. (A and B) Coronal sections of adult midbrain after 30 minutes of HRP (brown) circulation in wildtype (A) and *mfsd2aa* mutant fish (B). Wildtype adults confine the HRP within the blood vessels, but *mfsd2aa* mutants leak HRP into the brain parenchyma. The scale bar represents 200 µm. (C and D) TEM images of individual blood vessel cross-sections of adult wildtype (C) and *mfsd2aa* mutant fish (D). Endothelial cells (EC) are pseudo-colored green, pericytes (P) are pseudo-colored purple and red blood cells (RBC) are pseudo-colored red. (E and F) High magnification images of endothelial cells outlined with white dashed lines of wildtype (E) and *mfsd2aa* mutants (F). The images are oriented with the lumen (L) on top and the ablumen (A) on the bottom. *Mfsd2aa* mutants appear to have normal tight junctions (green arrowhead) but elevated levels of luminal and abluminal vesicles (magenta arrows). The scale bars represent 200 nm. (G and H) Quantification of the vesicular densities both on the luminal (G) and abluminal (H) side of the endothelial cells reveals that *mfsd2aa* mutants have increased vesicular densities. N=4 fish, each marked with a different color, with 15 blood vessels quantified for each fish. **** p < 0.0001, ** p < 0.01.

## Discussion

One of the major advantages to studying the BBB in zebrafish is the ability to perform live imaging of tracer permeability dynamics. We provide some of the first data on the dynamics of immature barrier leakage. At 3 dpf, we observed two types of leakage, the gradual overall increase in the parenchyma and rare transient bursts, both from a migrating sprout and established blood vessels. We attribute the gradual diffuse increase in parenchymal tracer uptake to the high levels of vesicular trafficking observed in our TEM analyses in which the endothelial basement membrane became filled with gold nanoparticles. Therefore, the parenchymal Dextran intensity measurements provide a direct proxy for the rates of BBB transcytosis *in vivo*. The rarer bursts of leakage, on the other hand, could be due to fleeting tight junctional ruptures between neighboring endothelial cells rather than increased vesicular trafficking. As these events were extremely scarce during our time lapse imaging, it would be nearly impossible to capture these potential junctional breaches by TEM to confirm this hypothesis. However, with continuing advances in the resolution of fluorescence microscopy, we could use zebrafish to resolve whether these two types of leakage, gradual and bursting, proceed through the same or different subcellular routes into the brain parenchyma, either paracellularly or transcellularly.

Among the various clathrin-independent transcytotic pathways (Tuma and Hubbard, 2003; Sandvig et al., 2018), caveolae are particularly abundant in vascular endothelial cells (Frank et al., 2003). Caveolae are caveolin-coated 50-100 nm flask-shaped invaginations of the plasma membrane (Palade, 1953; Palade, 1961). Furthermore, the suppression of caveolae-mediated transcytosis regulates the development and function of both the mouse blood-retinal barrier and the BBB (Andreone et al., 2017; Chow and Gu, 2017). Taken together, this suggests that the increased transcytosis during early larval stages is most likely caveolae mediated. Interestingly, the linear profile of Dextran uptake in the midbrain at 3 dpf closely resembles the caveolae-mediated uptake of a fluorescently-conjugated aminopeptidase P (APP) antibody in the mouse lung (Oh et al., 2007). While similarly linear, the scale is on the order of minutes in the larval zebrafish brain versus seconds in the mouse lung. This discrepancy in timing is most likely due to the large difference in the vesicular densities between the immature BBB endothelium (average of 2 vesicles/µm^2^) and the continuous endothelium of peripheral tissues, which ranges from 30 to 98 vesicles/µm^2^ on the luminal membrane of diaphragm and myocardial endothelium (Simionescu et al., 1974). Since loss of caveolin 1 is sufficient to precociously seal the blood-retinal barrier at the angiogenic front (Chow and Gu, 2017), it would be interesting in future work to examine whether zebrafish caveolin 1 mutants (Cao et al., 2016), which are viable as homozygotes, also exhibit an earlier onset of BBB maturation.

BBB permeability is tightly regulated by endothelial cell interactions with pericytes and astroglial cells. For the first time, we visualized their locations under TEM in zebrafish throughout barrier development. Pericytes have been shown to be essential for establishing the mammalian BBB (Armulik et al., 2010; Bell et al., 2010; Daneman et al., 2010). Similarly, pericyte deficient notch3^fh332^ fish display increased BBB permeability in a tight junction independent manner like pericyte-deficient mice (Wang et al., 2014). Our TEM data show that zebrafish pericytes are in close contact with brain endothelial cells (Figures 2 and 5), even at the earliest stage examined (3 dpf; Figure 2), and are embedded within the endothelial basement membrane as in mammals. Our subcellular localization data is in line with a growing body of evidence for the conserved role of pericytes in the zebrafish BBB (Wang et al., 2014; Lei et al., 2017). While zebrafish lack stellate astrocytes, they possess radial glia that express several astrocytic markers, such as Gfap, glutamine synthetase (GS), and Aqp4 (Jeong et al., 2008; Grupp et al., 2010). Previous studies have disagreed on the extent of radial glia interactions with BBB endothelial cells in zebrafish. Some studies report glia-BBB interactions that are similar to those observed in mammals (Jeong et al., 2008), while others report little to no interaction (Grupp et al., 2010). Our TEM data in adult zebrafish do not find a necessity for glial interactions with the brain vasculature, with endothelial cells occasionally in contact with electron-light glia (Figure 5B), but often not (Figure 5A). This appears to be a unique feature of the zebrafish BBB and should be considered carefully when using zebrafish as a model system for the BBB.

Taken together this study provides a thorough characterization of the development of the zebrafish BBB, highlighting regional differences in timing of maturation and capturing the dynamics of the immature BBB. Furthermore, our developmental TEM series provides the first direct evidence of vesicular trafficking regulating zebrafish BBB development. Finally, we have shown that this down-regulation of vesicular trafficking is necessary for BBB formation, as *mfsd2aa* mutants display increased barrier permeability due to unsuppressed transcytosis. We hope that this work will serve as a launching pad for future studies using zebrafish to understand the molecular regulators of BBB development and homeostasis in vertebrates.

## Author Contributions

N.M.O., S.G.M. and C.G. conceived the project and designed experiments. N.M.O. performed all experiments and analyzed all data. N.M.O., S.G.M. and C.G wrote the manuscript.

## Acknowledgments

We thank members of the Gu and Megason laboratories for data discussion and comments on the manuscript; Dr. Zach O’Brown for discussions and comments on the manuscript; Dr. Bela Anand-Apte (Cleveland Clinic) for providing the transgenic *l-fabp:DBP-EGFP* fish line (Xie et al., 2010) and Dr. Leonard Zon for providing the transgenic Tg(*kdrl:HRAS-mCherry*) line; and the HMS Electron Microscopy Core Facility, with special thanks to Louise Trakimas for all of her assistance in troubleshooting and preparing the TEM samples. This work was supported by the Damon Runyon Cancer Foundation (N.M.O.), the Mahoney postdoctoral fellowship (N.M.O.), the Fidelity Biosciences Research Initiative (C.G.), and the NIH DP1 NS092473 Pioneer Award (C.G.). The research of C.G. was also supported in part by a Faculty Scholar grant from the Howard Hughes Medical Institute.

## Materials and Methods

### Zebrafish Strains and Maintenance

Zebrafish were maintained at 28.5°C following standard protocols (Westerfield, 1993). All zebrafish work was approved by the Harvard Medical Area Standing Committee on Animals under protocol number 04487. Adult fish were maintained on a standard light-dark cycle from 9 am to 11 pm. Adult fish, age 3 months to 2 years, were crossed to produce embryos and larvae. These studies used the AB wildtype strain and the transgenic strains Tg(*l-fabp:DBP-EGFP)*, Tg(*kdrl:mCherry*), and Tg(*kdrl:HRAS-mCherry*).

### Tracer Injections

Larvae were immobilized with tricaine and placed in an agarose injection mold with their hearts facing upwards. 2.3 nl of Alexa Fluor 405 NHS Ester (Thermo Fisher: A30000) or Alexa Fluor 647 10 kDa Dextran (Thermo Fisher:D22914) fluorescently conjugated tracers (10 mg/ml) were injected into the cardiac sac using Nanoject II (Drummond Scientific, Broomall, PA). Embryos were then mounted with 1.5% low gelling agarose (Sigma: A9414) in embryo water on 0.17 mm coverslips and imaged live within 2 hours post injection. For developmental electron microscopy experiments, 2.3 nl of 5 nm NHS-activated gold nanoparticles (Cytodiagnostics: CGN5K-5-1, ∼1.1^14^ particles/ml in PBS) were injected into the cardiac sac just as for the fluorescently conjugated tracers. After 5 minutes of circulation, the fish were fixed for electron microscopy. Adults were briefly immobilized with tricaine and retroorbitally injected with 3 µl of HRP (2 mg/ml dissolved in PBS) using a 10 µl Hamilton syringe. After 30 minutes of HRP circulation, the brains were fixed in 4% PFA overnight at 4C. Following fixation the brains were washed three times and sectioned coronally with a vibratome (50 µm). Sections were then stained at room temperature in 0.05 M Tris-HCl pH 7.6 buffer containing 0.5 mg/ml 3-39 diaminobenzidine (DAB, Sigma Aldrich) and 0.01% hydrogen peroxide.

### Transmission Electron Microscopy (TEM)

Fish were anesthetized with tricaine and initially fixed by immersion in 4% paraformaldehyde (VWR:15713-S) /0.1M sodium-cacodylate (VWR:11653). Following this initial fixation, the larval fish and adults with exposed brains were further fixed for 7-14 days in 2% glutaraldehyde (Electron Microscopy Sciences: 16320)/ 4% paraformaldehyde/ 0.1M sodium-cacodylate at room temperature. Following fixation, larvae or dissected brains were washed overnight in 0.1M sodium-cacodylate. Entire larval heads or coronal vibratome free-floating sections of adult brains (50 µm) were post-fixed in 1% osmium tetroxide and 1.5% potassium ferrocyanide, dehydrated, and embedded in epoxy resin. Ultrathin sections of 80 nm were then cut from the block surface and collected on copper grids. The adult sections were counter-stained with Reynold’s lead citrate prior to imaging.

### CRISPR Mutants

*Mfsd2aa* mutant fish were generated by injection of Cas9 RNA and a single guide RNA (5’- GGTGTGTTTTGCGATCGGAG-3’) targeting exon 3 into single-cell fertilized wildtype embryos. *Mfsd2ab* mutant fish were generated by injection of Cas9 protein and a single guide RNA (5’- TGAGAGCAGAGTAGGGCACG-3’) targeting exon 5 into single-cell fertilized double transgenic Tg(*l-fabp:DBP-EGFP; kdrl:mCherry)* embryos. F0 injected fish were raised, outcrossed to wildtype fish and screened for potential mutant founders by PCR and sequencing. The stable *mfsd2aa* mutant line was genotyped using 5’-AAATCACCTCTTCCAGTGAGGA-3’ and 5’- ATAGTAACAAAACGATGCTGAGCC-3’ primers. *Mfsd2ab* mutants were genotyped using 5’- GTCTACTCCATTTGCTGTACTTTGC-3’ and 5’-CAGGTCAATCTCAGTGCTGATACAG-3’ primers.

### Imaging

All live imaging of tracer permeability was performed on a Leica SP8 laser scanning confocal microscope. Time lapse imaging was performed on the SP8 using a resonance scanner. A 1200EX electron microscope (JOEL) equipped with a 2k CCD digital camera (AMT) was used for all TEM studies. Images were visualized and quantified using ImageJ (NIH) and Adobe Photoshop. Time lapse videos were visualized as 3D reconstructions and cropped to highlight particular blood vessels of interest using Imaris software (Bitplane).

### Tracer Permeability Quantification

All quantification was performed on blinded image sets. Parenchymal cells containing injected tracers outside of the blood vessels were manually counted throughout z-stacks that spanned the entire larval brain in depth. Z-stacks were collected with 0.59 µm z-steps using a 25x water immersion objective on a Leica SP8 laser scanning confocal microscope. The Dextran intensity was measured in six parenchymal regions of average intensity projections of the time lapse videos and averaged as a single value per fish. These values were normalized to the average fluorescence intensity in the lumen at each time point.

### TEM Quantification

For all TEM quantifications, vesicular density values were calculated from the number of non-clathrin coated flask-shaped vesicles per µm of endothelial luminal or abluminal membrane for each image collected. Embryonic basement membrane gold intensity was measured in three regions per endothelial cell and normalized to luminal gold intensity values within individual images using ImageJ. All images for analysis were collected at 12000x magnification on the JOEL 1200EX electron microscope. 10-15 vessels were quantified for each fish, with each color representing a different fish.

### Statistical Analysis

All statistical analyses were performed using Prism 8 (GraphPad Software). Two group comparisons were analyzed using an unpaired two-tailed t test. Nested t tests were employed for all electron microscopy comparisons to account for the actual N versus vessels analyzed. Multiple group comparisons were analyzed with one-way ANOVA, followed by a post hoc Tukey’s multiple comparison test. The time lapse leakage dynamics were analyzed with a Mann-Whitney U test to discriminate whether the two patterns of leakage accumulation were different. Sample size for all experiments was determined empirically using standards generally employed by the field, and no data was excluded when performing statistical analysis. Standard error of the mean was calculated for all experiments and displayed as errors bars in graphs. Statistical details for specific experiments, including exact n values and what n represents, precision measures, statistical tests used, and definitions of significance can be found in the Figure Legends.

---

**Figure 1 – Supplement 1.**
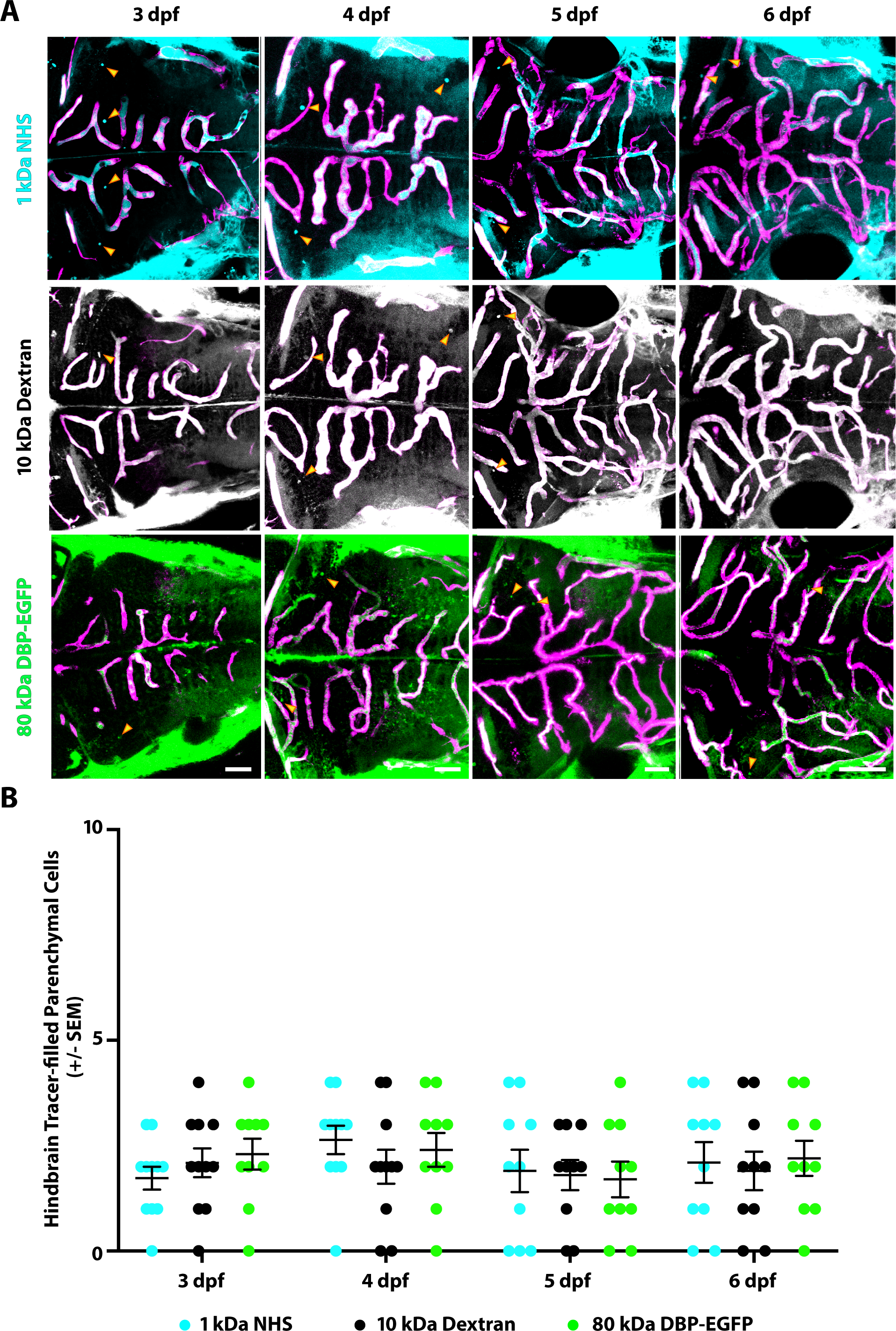
The hindbrain has a functional BBB at 3 dpf. (A) Representative dorsal view maximum intensity projections of larval zebrafish hindbrains at different developmental stages reveals low permeability at 3 dpf that stays constant until 6 dpf. This low permeability was observed with two injected tracers of different sizes, a 1 kDa NHS (turquoise) and a 10 kDa Dextran (white), as well with an 80 kDa transgenic serum protein DBP-EGFP (green). Yellow arrowheads demarcate the few tracer-filled parenchymal cells outside of the vasculature (magenta). Scale bars represent 50 µm. (B) Quantification of tracer-filled parenchymal cells in the hindbrain between 3 and 6 dpf reveals low tracer uptake beginning at 3 dpf. There was no difference in the number of parenchymal cells that picked up the different tracers at any time point. N = 10-11 fish, each represented as a single dot on the plot. The mean and the standard error are drawn in black for each tracer and stage.

**Figure 4 – Supplement 1.**
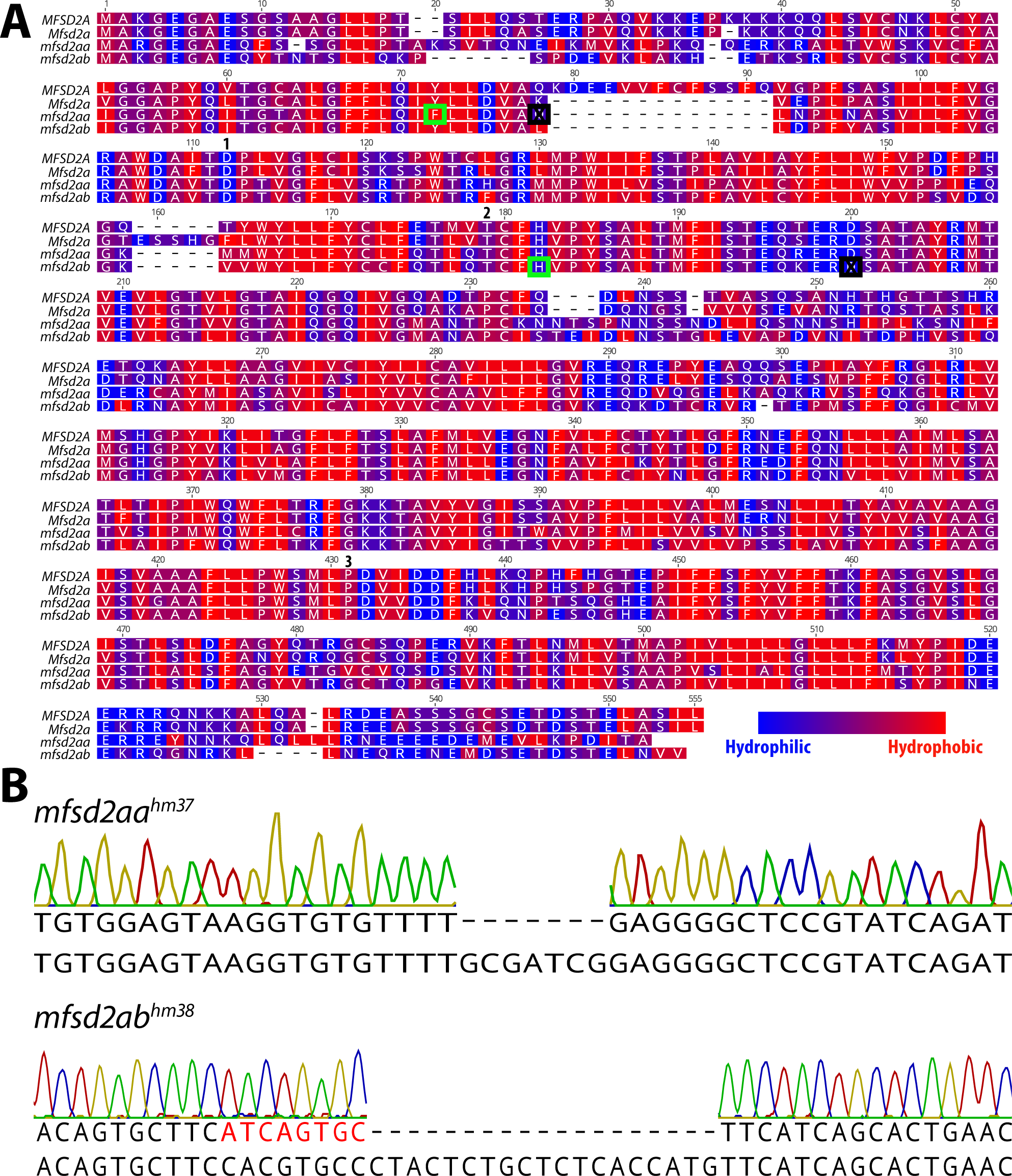
Zebrafish have 2 *Mfsd2a* paralogues, *mfsd2aa* and *mfsd2ab*. (A) Protein alignment of human MFSD2A, mouse Mfsd2a, zebrafish mfsd2aa, and zebrafish mfsd2ab illustrated with a hydrophobicity scale. Red amino acids are hydrophobic and blue amino acids are hydrophilic. Both zebrafish paralogues are highly similar to the human and mouse proteins. Green boxes highlight the genetic lesions in (B) and the black boxes mark the predicted premature stop codons caused by these mutations in *mfsd2aa^hm37/hm37^* and *mfsd2ab^hm38/hm38^*mutants. Numbers mark amino acid residues with mutations that have been shown to impact Mfsd2a function:1. Andreone et al., 2017, 2. Guemez-Gamboa et al., 2015, 3. Harel et al., 2018. (B) Sanger sequencing of the *mfsd2aa^hm37^* and *mfsd2ab^hm38^*mutations. *Mfsd2aa^hm37/37^* mutants have a 7 base pair deletion in exon 2 that is predicted to lead to a premature stop codon at amino acid 82 (A; black box). *Mfsd2ab^hm38/hm38^*mutants have an 8 base pair insertion (red letters) and a 19 base pair deletion in exon 5 that is predicted to lead to a premature stop codon at amino acid 175 (A; black box).

**Figure 4 – Supplement 2.**
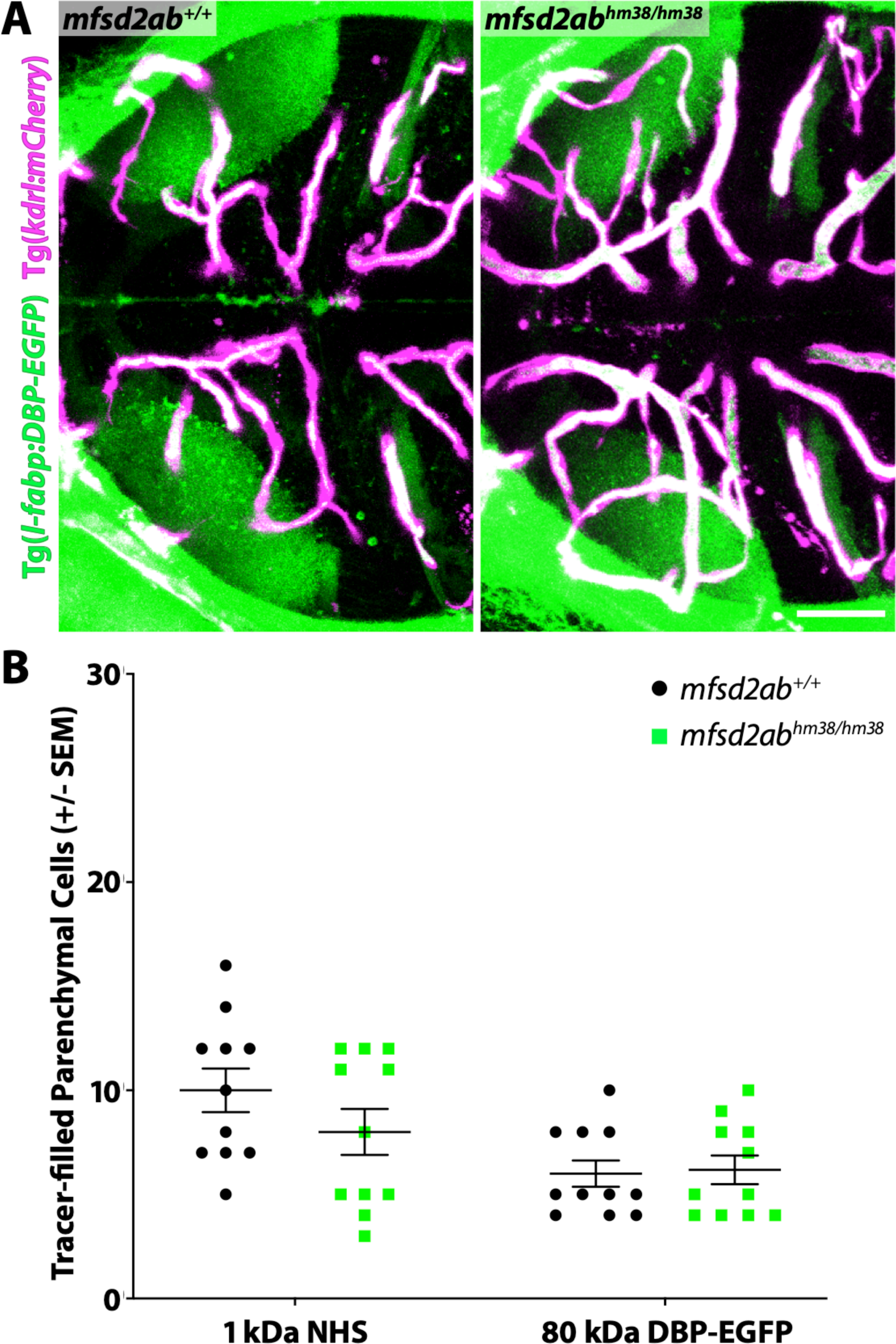
*Mfsd2ab* mutants do not have altered BBB permeability. (A) Representative maximum intensity projection images of the midbrain of wildtype and *mfsd2ab* mutants expressing the fluorescently labelled 80 kDa transgenic serum protein DBP-EGFP (green) at 5 dpf. *Mfsd2ab* mutants have a similarly low number of DBP-EGFP-filled parenchymal cells compared to wildtype siblings. The scale bar represents 50 µm. (B) Quantification of tracer-filled (NHS and DBP-EGFP) parenchymal cells in the midbrain of wildtype (black) and *mfsd2ab* mutants (green). *Mfsd2ab* mutants display no significant barrier permeability defects, both for the injected NHS and the endogenous transgene DBP-EGFP (A). Each individual fish measured is displayed as a single point. The mean and the standard error are drawn in black.

**Figure 4 – Supplement 2.**
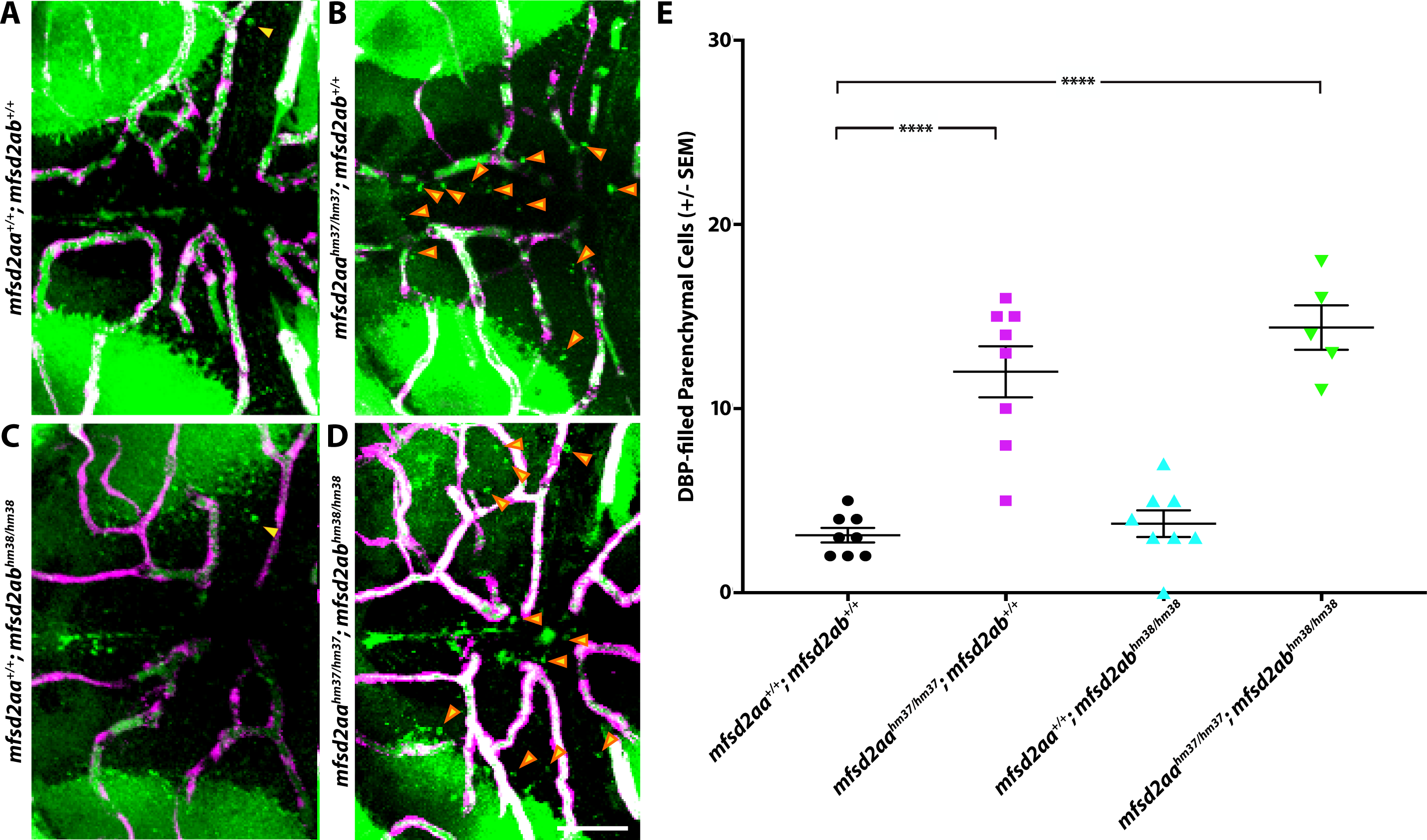
*Mfsd2aa^hm37/hm37^; mfsd2ab^hm38/hm38^*double mutants display similar increased BBB permeability to *mfsd2aa^hm37/hm37^* fish. (A-D) Representative maximum intensity projection images of the midbrain of wildtype (A; *mfsd2aa^+/+^; mfsd2ab^+/+^*), *mfsd2aa^hm37/hm37^*single mutants (B), *mfsd2ab^hm38/hm38^*single mutants (C), and *mfsd2aa^hm37/hm37^; mfsd2ab^hm38/hm38^*double mutants (D) expressing the fluorescently labelled 80 kDa transgenic serum protein DBP-EGFP (green) at 5 dpf. *Mfsd2aa^hm37/hm37^; mfsd2ab^hm38/hm38^* double mutants display a similar level of increased tracer-filled parenchymal cells (yellow arrowheads) to the *mfsd2aa^hm37/hm37^*single mutants. The scale bar represents 50 µm. (E) Quantification of DBP-EGFP-filled parenchymal cells in the midbrain of wildtype (black), *mfsd2aa^hm37/hm37^* mutants (magenta), *mfsd2ab^hm38/hm38^* mutants (turquoise), and *mfsd2aa^hm37/hm37^; mfsd2ab^hm38/hm38^* double mutants (green). Each individual fish measured is displayed as a single point. The mean and the standard error are drawn in black. **** p < 0.0001.

**Figure 5 – Supplement 1.**
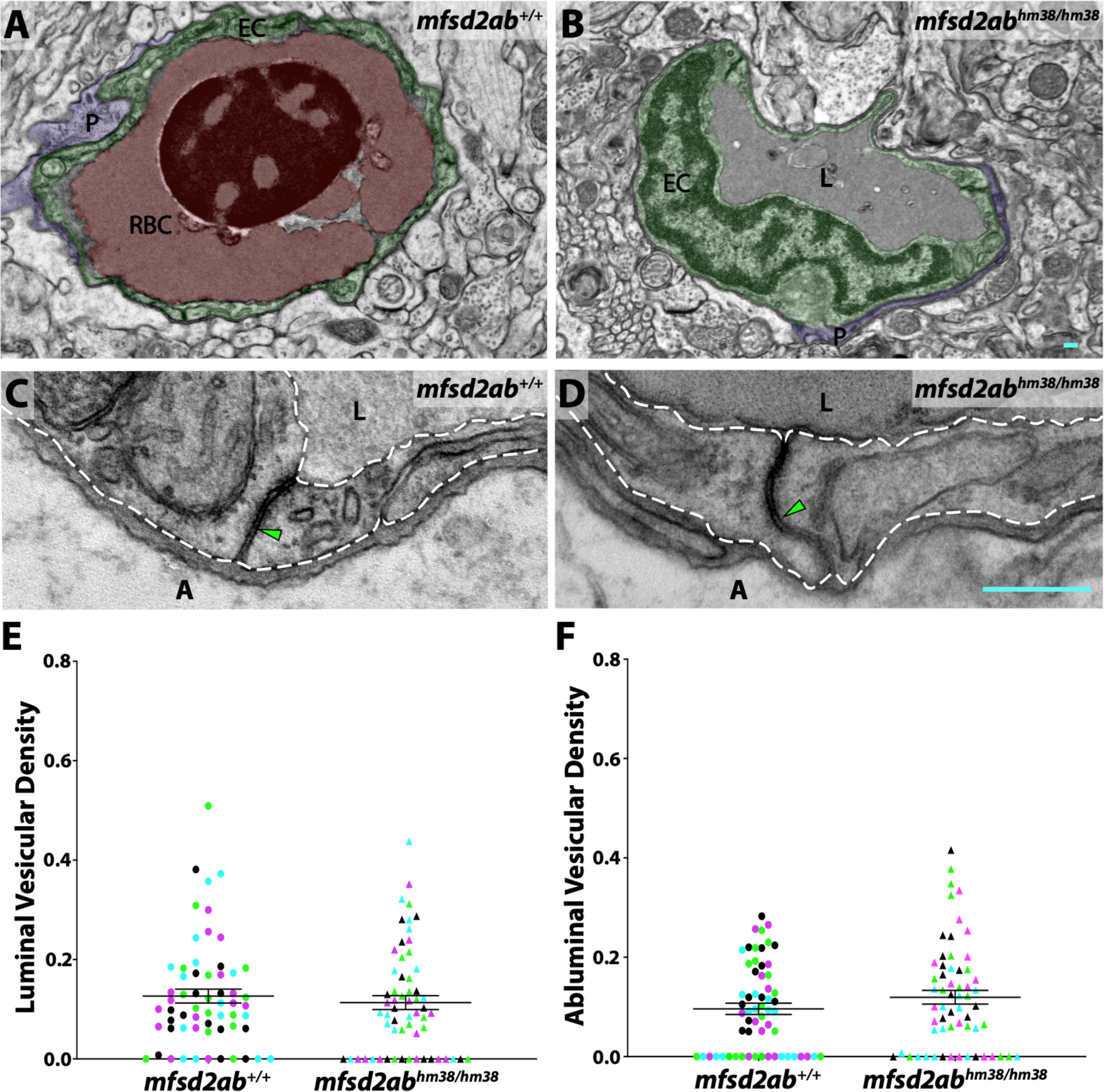
M*fsd2ab* mutants exhibit normal vascular maturation. (A and B) TEM images of individual blood vessel cross-sections of adult wildtype (A) and *mfsd2ab* mutants (B). Endothelial cells (EC) are pseudo-colored green, pericytes (P) are pseudo-colored purple and red blood cells (RBC) are pseudo-colored red when present in the lumen (L). (C and D) High magnification images of endothelial cells outlined with white dashed lines of wildtype (C) and *mfsd2ab* mutants (D). The images are oriented with the lumen (L) on top and the ablumen (A) on the bottom. *Mfsd2ab* mutants appear to have normal tight junctions (green arrowhead) and levels of luminal and abluminal vesicles. The scale bars represent 200 nm. (E and F) Quantification of the vesicular densities both on the luminal (E) and abluminal (F) side of the endothelial cells reveals that *mfsd2ab* mutants have similar vesicular densities to wildtype siblings. N=4 fish, each marked with a different color, with 15 blood vessels quantified for each fish.

